# Gene discoveries in autism are biased towards comorbidity with intellectual disability

**DOI:** 10.1101/715755

**Authors:** Matthew Jensen, Corrine Smolen, Santhosh Girirajan

## Abstract

Autism typically presents with a highly heterogeneous set of features, including frequent comorbidity with intellectual disability (ID). The overlap between these two phenotypes has confounded the accurate diagnosis and discovery of genetic factors associated with autism. We analyzed genetic variants in 2,290 individuals with autism from the Simons Simplex Collection (SSC) who have either ID or normal cognitive function to determine whether genes associated with autism also contribute towards ID comorbidity. We found that individuals who carried variants in a set of 173 reported autism-associated genes showed decreased IQ (p=5.49×10^−6^) and increased autism severity (p=0.013) compared with individuals without such variants. A subset of autism-associated genes also showed strong evidence for ID comorbidity in published case reports. We also found that individuals with high-functioning autism (IQ>100) had lower frequencies of CNVs (p=0.065) and LGD variants (p=0.021) compared with individuals who manifested both autism and ID (IQ<70). These data indicated that *de novo* LGD variants conferred a 1.53-fold higher risk (p=0.035) towards comorbid ID, while LGD mutations specifically disrupting autism-associated genes conferred a 4.85-fold increased risk (p=0.011) for comorbid ID. Furthermore, *de novo* LGD variants in individuals with high-functioning autism were more likely to disrupt genes with little functional relevance towards neurodevelopment, as demonstrated by evidence from pathogenicity metrics, expression patterns in the developing brain, and mouse model phenotypes. Overall, our data suggest that *de novo* pathogenic variants disrupting genes associated with autism contribute towards autism and ID comorbidity, while other genetic factors are likely to be causal for high-functioning autism.

---

Autism spectrum disorder, which presents in children with social communication difficulties, repetitive behavior, and restricted interests^1^, is a highly heterogeneous neurodevelopmental disorder characterized by complex genetic etiology and strong comorbidity with other developmental disorders^2^. For example, approximately 30% of individuals with autism also manifest with intellectual disability (ID)^3^, defined^1^ by IQ scores <70. The high degree of cooccurrence of autism with ID has been shown to confound accurate diagnosis of autism. In fact, we recently showed that 69% of individuals diagnosed with ID are likely to be recategorized and diagnosed with autism^4^. The diagnostic overlap between autism and ID suggests that *de novo* gene disruptive variants and copy-number variants (CNVs) identified in individuals ascertained for autism in large-scale studies could also be confounded by ID comorbidity. Here, using genetic and phenotypic data from 2,290 individuals with autism from the Simons Simplex Collection (SSC)^5^, we show that gene discoveries in autism are biased towards genes that contribute towards both autism and comorbid ID.

We analyzed rare *de novo* likely-gene disruptive (LGD) variants from exome sequencing data^6,7^, disease-associated copy-number variants (CNVs) from microarrays^8^, and Full-scale IQ and Social Responsiveness Scale (SRS) T-scores for SSC probands that were obtained from the Simons Foundation Autism Research Initiative^5^. As these data were de-identified, they were exempt from IRB review and conformed to the Helsinki Declaration. We first compared the phenotypes of 288 individuals with *de novo* LGD variants and 81 individuals with pathogenic CNVs to 1,921 individuals without such variants obtained from the SSC cohort. Similar to previous autism studies that identified correlations between *de novo* variants and IQ scores^9–12^, we found that individuals with *de novo* LGD variants (IQ=77.7, p=0.031, two-tailed Mann-Whitney test) or pathogenic CNVs (IQ=76.3, p=0.002) had a significant decrease in IQ scores compared with individuals without such variants (IQ=82.3) (**Figure 1A**). However, no differences in autism severity, measured using SRS T-scores, were observed between groups of individuals with and without pathogenic variants (p=0.104 for LGD variants and 0.963 for CNVs) (**Figure 1A**). This suggests that pathogenic variants in general contribute to ID independent of autism severity, although this could also be due to an ascertainment bias in the SSC cohort towards individuals with severe autism.

**Figure 1.**
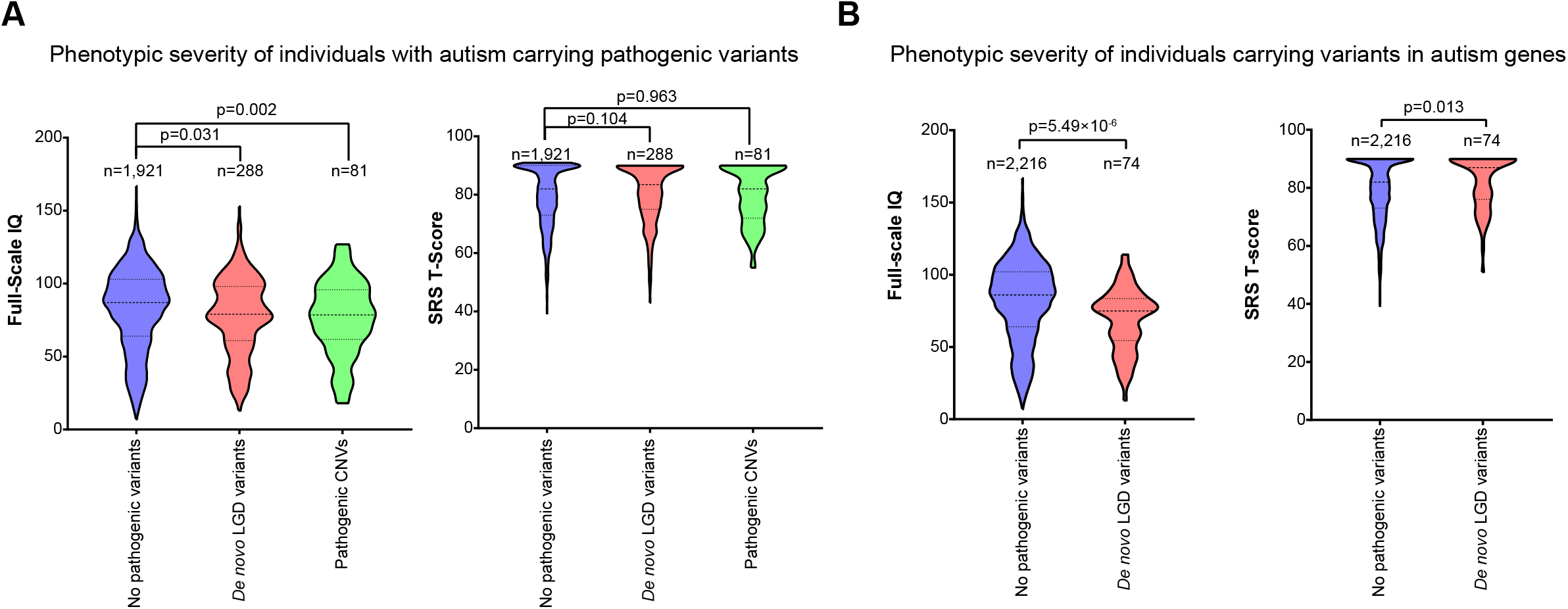
Phenotypic comparison of individuals with autism from the SSC cohort with and without pathogenic variants. **(A)** Individuals with pathogenic variants (*de novo* LGD and CNV) had a significantly lower IQ than individuals without pathogenic variants, but no change in autism severity (SRS T-score) was observed between the three groups. **(B)** Individuals with *de novo* LGD variants in candidate autism genes had a lower IQ and more severe autism phenotypes than individuals without such variants. n indicates sample size, p-values were derived from two-tailed Mann-Whitney tests, and dotted lines within each plot indicate the median and first and third quartiles. All statistics were calculated using R v.3.4.2 (R Foundation for Statistical Computing, Vienna, Austria).

We further identified individuals carrying *de novo* LGD variants in 173 autism-associated genes, defined as genes with recurrent *de novo* variants reported in multiple databases of sequencing studies (**Table S1**). These genes included tier 1 genes (>2 *de novo* LGD variants) from the Developing Brain Disorders Gene Database^13^, genes with >5 non-SSC *de novo* LGD variants from denovo-db^14^, and SFARI Gene tiers 1 and 2 (https://gene.sfari.org/). We found that individuals carrying *de novo* LGD variants in autism-associated genes had decreased IQ (n=74, IQ=69.1, p=5.49×10^−6^, two-tailed Mann-Whitney test) and increased SRS T-scores (SRS=82.4, p=0.013) compared with individuals without LGD variants (n=2,216, IQ=81.9, SRS=79.6), implying that candidate autism genes contribute to both autism and ID phenotypes (**Figure 1B**). To validate this finding, we examined 76 published case reports of affected individuals with pathogenic variants in a subset of 22 autism genes that appeared in all three autism gene databases (**Table 1, Table S2**). For example, recent case studies have identified autism cooccurring with ID in 21 individuals with *de novo SHANK3* variants^15^, 19 individuals with *NRXN1* variants^16^, and 18 individuals with *TCF20* variants^17^. Overall, 460/497 (92.6%) individuals with autism described in these studies had ID features, emphasizing that variants in these genes contribute to a severe form of autism with comorbid ID **(Table 1)**.

**Table 1.** Individuals carrying variants in autism-associated genes with comorbid ID.

| **Autism-associated genes** | **Cases with ID** | **Autism cases with or without ID** | **Autism cases with comorbid ID** |
| --- | --- | --- | --- |
| *ADNP* | 134/134 | 114/134 | 114/114 |
| *ANK2* | 1/1 | 1/1 | 1/1 |
| *ANKRD11* | 10/10 | 9/10 | 9/9 |
| *ARID1B* | 137/153 | 80/153 | 80/80 |
| *ASH1L* | 14/14 | 4/14 | 4/4 |
| *ASXL3* | 18/19 | 16/19 | 15/16 |
| *BCL11A* | 11/16 | 4/16 | 3/4 |
| *CHD2* | 3/3 | 3/3 | 3/3 |
| *CHD8* | 56/75 | 61/75 | 46/61 |
| *CUL3* | 1/1 | 1/1 | 1/1 |
| *DDX3X* | 97/97 | 33/97 | 33/33 |
| *DYRK1A* | 7/26 | 9/26 | 7/9 |
| *KMT2A* | 76/99 | 12/99 | 10/12 |
| *MECP2* | 1/1 | 1/1 | 1/1 |
| *MYT1L* | 1/1 | 1/1 | 1/1 |
| *NRXN1* | 45/60 | 34/60 | 23/34 |
| *POGZ* | 48/49 | 29/49 | 29/29 |
| *SCN2A* | 19/33 | 9/33 | 7/9 |
| *SETD5* | 15/16 | 7/16 | 7/7 |
| *SHANK3* | 35/37 | 26/37 | 24/26 |
| *SYNGAP1* | 11/23 | 11/23 | 11/11 |
| *TCF20* | 47/48 | 32/48 | 31/32 |
| **Total** | **787/916 (85.9%)** | **497/916 (54.3%)** | **460/497 (92.6%)** |

We next compared genetic data from 397 SSC individuals (17.3% of the SSC cohort) with “high-functioning autism”, defined as having severe autism and average or above-average IQ scores (SRS>75 and IQ>100), to 562 individuals (24.5%) with both autism and ID (SRS>75 and IQ<70). Individuals with high-functioning autism had a significantly lower (p=0.021, one-tailed Fisher’s Exact test) frequency of *de novo* LGD variants (42/397, 10.6%) than individuals with autism and ID (86/562, 15.3%). Similarly, individuals with high-functioning autism were less likely (p=0.065) to carry pathogenic CNVs (9/397, 2.3%) than individuals with autism and ID (24/562, 4.3%). In fact, *de novo* LGD variants conferred a 1.53-fold higher likelihood of manifesting ID among individuals with autism (p=0.035, 95% confidence interval 1.03-2.26), and pathogenic CNVs similarly conferred a 1.92-fold increased risk for co-occurrence of ID among individuals with autism (p=0.099, 95% CI 0.88-4.18). We replicated these observations by analyzing an additional combined cohort of 2,357 individuals from both the SSC and the Autism Sequencing Collection^18^. Here, individuals with autism and ID had a significantly higher rate (p=3.04×10^−6^, one-tailed Student’s t-test) of *de novo* variants in genes intolerant to variation, as measured by probability of Loss-of-function Intolerant (pLI) score >0.9 (70/643, 10.8%), than individuals manifesting autism but not ID (114/1747, 6.65%). We also found that only 3/397 (0.8%) individuals in the SSC cohort with high-functioning autism carried *de novo* LGD variants in autism-associated genes, including *ANK2, HIVEP3,* and *BAZ2B*. This frequency was not significantly different from the expected frequency of *de novo* variants in the general population (p=0.095, one-tailed Student’s T-test), as calculated from gene-specific probabilities of *de novo* nonsense and frameshift variants from a sequence context-dependent model^9^. In contrast, 20/562 (3.6%) individuals with autism comorbid with ID carried *de novo* LGD variants in autism-associated genes, such as *CHD8*, *SCN2A*, and *SYNGAP1*, representing a 19.2-fold enrichment of variants compared with the expected rate in the general population (p=9.48×10^−6^). Thus, *de novo* LGD variants in autism genes conferred a 4.85-fold increased risk (p=0.011, 95% CI 1.43-16.42) towards comorbid ID in individuals with autism.

We further sought to determine the biological relevance of the 42 genes with *de novo* LGD variants identified in individuals with high-functioning autism, and found that these genes in aggregate had less functional relevance towards neurodevelopment than the reported autism-associated genes. For example, genes with *de novo* LGD variants in individuals with high-functioning autism were less resistant to genetic variation than reported autism-associated genes, as measured by Residual Variation Intolerance Score (RVIS) (p=4.00×10^−4^, Mann-Whitney two-tailed test) and pLI percentile (p=9.77×10^−7^) gene metrics^19,20^ (**Figure 2A).** In fact, while the RVIS and pLI percentiles of the reported autism genes were clustered below the thresholds for pathogenicity (RVIS <20^th^ percentile and pLI <18^th^ percentile, or raw score >0.9), genes disrupted among individuals with high-functioning autism were evenly distributed across the range of percentiles. Additionally, we tested the enrichment of each gene set for specific expression in brain regions during development, based on expression data derived from the BrainSpan Atlas^21^, using the Specific-Expression Analysis (SEA) online tool^22^. While autism genes were enriched for specific expression in the cortex (p=3.13×10^−4^, Fisher’s Exact test with Benjamini-Hochsberg correction) and cerebellum (p=0.020) during early fetal development^22^, genes with *de novo* LGD variants in high-functioning autism individuals were not enriched for any specific expression patterns in the developing brain (**Figure 2B**). Furthermore, mouse models of genes identified in individuals with high-functioning autism, whose phenotypic data were obtained from the Mouse Genome Informatics database^24^, were significantly less likely to manifest nervous system (p=4.90×10^−3^, one-tailed Fisher’s Exact test with Benjamini-Hochsberg correction) and behavioral/neurological (p=0.037) phenotypes than mouse models of reported autism-associated genes (**Figure 2C**). These findings suggest that genes with *de novo* LGD variants in individuals with high-functioning autism are less pathogenic in humans and model organisms, and therefore may not necessarily contribute towards the specific high-functioning autism phenotype.

**Figure 2.**
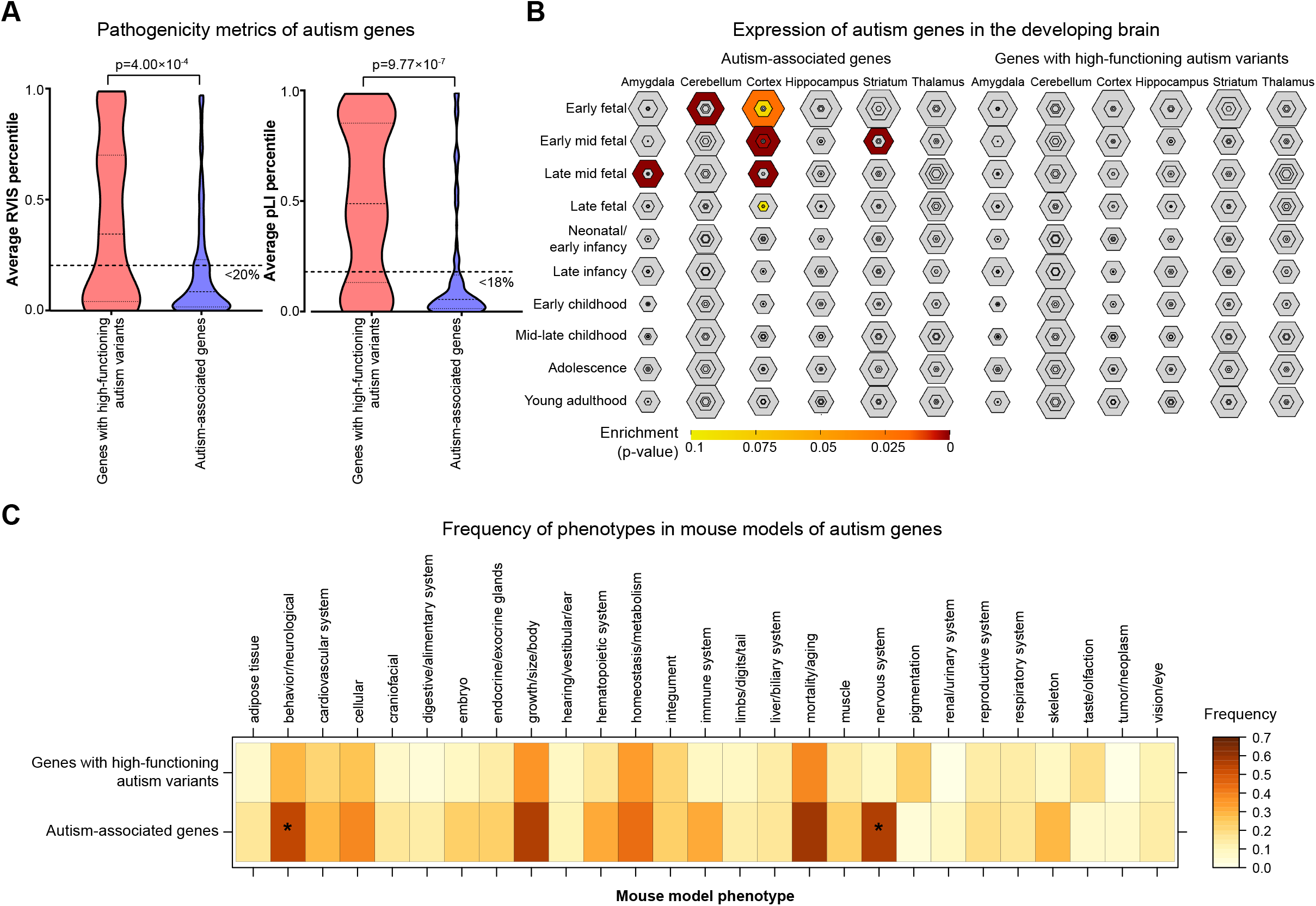
Functional analysis of genes with *de novo* LGD variants in individuals with high-functioning autism. (A) Genes with *de novo* LGD variants in individuals with high-functioning autism had lower average RVIS (left) and pLI (right) percentile scores than those for reported autism-associated genes. Thick dotted lines across the violin plots indicate thresholds for gene pathogenicity: <20^th^ percentile for RVIS and <18^th^ percentile for pLI (>0.9 raw score). Thin lines within the violin plots indicate the median and first and third quartiles. p-values were derived from two-tailed Mann-Whitney tests. (B) Expression of genes with *de novo* variants in individuals with high-functioning autism and autism-associated genes in the developing human brain. Autism-associated genes were enriched for specific expression in the cortex and cerebellum during early development, while no enrichment was seen in the genes identified in individuals with high-function autism. Hexagon sizes represent the number of genes preferentially expressed in each brain tissue and timepoint, while colors of the hexagons represents p-values for the enrichment of autism genes among each set of preferentially-expressed genes. (C) Frequency of phenotypes observed in mouse knockout models for genes with *de novo* LGD variants in individuals with high-functioning autism compared with reported autism-associated genes. * indicates p<0.05 with Benjamini-Hochsberg correction.

Our data indicate that pathogenic variants such as *de novo* LGD variants and CNVs contribute to autism phenotypes primarily in individuals with comorbid ID, especially if the variants disrupt a gene previously associated with autism. Several themes regarding the study of high-functioning autism have emerged from these findings. *First*, the consistently high degree of comorbidity between autism and ID has led to an ascertainment bias towards individuals who manifest both disorders in large-scale sequencing cohorts, as it is difficult to exclude all individuals with comorbid disorders and still have adequate power to identify recurrent variants. Indeed, more than 80% of the SSC cohort had an IQ score less than 100, and the average IQ of the cohort (81.5) was 18.5 points below the population average. This bias has contributed to the identification of genes and CNV regions related to both autism and ID, as evidenced by the decreased IQ among carriers of variants in these genes as well as a high incidence of comorbid phenotypes reported in published case studies. Large-scale sequencing studies still hold a high value in uncovering shared biological mechanisms that could underlie both disorders^23^. However, understanding the biology of the core autism phenotypes would require concerted efforts to recruit individuals who specifically manifest high-functioning autism without ID.

*Second*, individuals with high-functioning autism are less likely to carry *de novo* LGD variants in candidate autism genes, and *de novo* variants in individuals with high-functioning autism tend to disrupt genes with less functional relevance towards neurodevelopment. These genes likely carry non-recurrent variants that either confer a small effect size towards autism risk on their own, or are not associated at all with neurodevelopment. We therefore propose that multiple genomic factors with varying effect sizes, such as missense variants, common variants, variants in regulatory and non-coding regions, or the combinatorial effects of inherited variants, contribute towards autism phenotypes without ID. For example, Schaaf and colleagues performed targeted sequencing of 21 candidate autism genes in 339 individuals with high-functioning autism^25^. They found that 2% of individuals carried *de novo* missense variants in candidate autism genes, such as *PTEN* and *FOXP2,* suggesting that allelic variants of differing severity within the same gene might contribute to distinct neurodevelopmental trajectories. Interestingly, the same study also found that 7% of individuals with high-functioning autism carried multiple inherited missense variants in candidate autism genes, potentially contributing to an oligogenic model for high-functioning autism phenotypes. Similarly, common variants have been found to contribute towards increased autism risk in individuals without ID^26,27^. For example, Grove and colleagues recently reported that the heritability attributed to common variants, including those primarily associated with cognitive ability and educational attainment, was three times lower in individuals with autism and ID compared with those without ID^27^. Finally, variants that may not contribute directly towards autism phenotypes themselves, including the *de novo* LGD variants observed in individuals with high-functioning autism, could still be responsible for subtler modification of the severity of autism or ID phenotypes.

Overall, our results emphasize the importance of dissecting phenotypic heterogeneity in family-based sequencing studies of complex diseases, especially those with a high frequency of comorbid disorders. While a larger cohort of individuals recruited specifically for high-functioning autism could identify associations with recurrent genes or different types of variants, these findings should be validated using functional studies to more fully differentiate the genetic causes for high-functioning autism from those for autism with comorbid ID.

## Supplemental data

Supplemental data include two supplemental tables in Excel file format.

## Acknowledgements

This work was supported by NIH R01-GM121907, SFARI Pilot Grant (#399894) and resources from the Huck Institutes of the Life Sciences to S.G., and NIH T32-GM102057 to M.J. The authors thank Fereydoun Hormozdiari (UC Davis), Lucilla Pizzo (Penn State), and Vijay Kumar (Penn State) for their helpful discussions and comments on the manuscript. We are grateful to all of the families at the participating Simons Simplex Collection (SSC) sites, as well as the principal investigators (A. Beaudet, R. Bernier, J. Constantino, E. Cook, E. Fombonne, D. Geschwind, R. Goin-Kochel, E. Hanson, D. Grice, A. Klin, D. Ledbetter, C. Lord, C. Martin, D. Martin, R. Maxim, J. Miles, O. Ousley, K. Pelphrey, B. Peterson, J. Piggot, C. Saulnier, M. State, W. Stone, J. Sutcliffe, C. Walsh, Z. Warren, E. Wijsman). We appreciate obtaining access to phenotypic data on the Simons Foundation Autism Research Initiative (SFARI) Base. Approved researchers can obtain the SSC data sets described in this study by applying at https://www.base.sfari.org.

## Author Contributions

M.J. and S.G. conceptualized the study, and M.J. and C.S. analyzed the data. M.J. and S.G. wrote the manuscript with input from all authors.

